# Evaluation of the effect of hydrocortisone in 2D and 3D HEp-2 cell culture

**DOI:** 10.1101/2021.03.26.437247

**Authors:** M.O. Fonseca, B.H. Godoi, N.S Da Silva, C. Pacheco-Soares

## Abstract

Cancer is one of the diseases with the highest incidence globally and that associated with the patient’s emotional state, can act positively or negatively in the treatment. Cortisol is a principal primary stress hormone in the human body. The corticoids can increase cell proliferation and reactive oxygen species that contribute to DNA damage. Prolonged exposure to stress can contribute to tissues becoming insensitive to cortisol, the primary human stress hormone. This study explores cortisol’s influence on tumor cell development, particularly in human cells of carcinoma of the human laryngeal (HEp-2). HEp-2 cells were exposed to increasing cortisol (hydrocortisone) concentrations for 24 or 48 hours, and cytotoxicity (MTT assay) proliferation assay (crystal violet assay), and immunolabeled 3D culture for fibronectin and FAK were analyzed. The group treated with hydrocortisone showed a significant increase in mitochondrial activity, as for the evaluation by the violet crystal, the treated group showed similar behavior to the control. The 3D culture showed dispersed cells within 24 hours with reduced FAK labeling; however, no changes were observed within 48 hours. Although some cases favored corticosteroid use in cancer patients, a more detailed analysis is necessary before prescribing them.

## 1. Introduction

Cancer is one of the most feared diseases of the 20th century and spreads with increasing incidence in the 21st century (Roy and Saikia, 2016). As regards occurrences of cancer, several factors do have identified as contributing factors, including genetic predisposition, exposure to environmental risk factors, contagion by certain viruses, cigarette use, and the ingestion of carcinogens ((Blackadar 2016; Chae and Lee 2019) (Seyfried, Shelton, and Mukherjee 2010).

Psychological factors can contribute to cancer development given the effects of emotional states on hormonal modification and alter the immune system stress influenced (Reiche, Nunes, and Morimoto 2004; Bedillion, Ansell, and Thomas 2019). From these possibilities, we increasingly find studies that seek to relate or measure possible influences of psychological and social aspects in the development and potential aggravation of oncological pathologies.

The disease’s symptoms and the related characteristics of the different treatment forms used to fight cancer significantly interfere with patients’ routines and quality of life, characterizing significant stressors in many cases (Bedillion, Ansell, and Thomas 2019; Spiegel and Giese-Davis 2003).

Studies relate factors associated with stress to tumor biology; those that describe the effects of glucocorticoids (GC) and their relationship with tumor cells’ biology and out (Vitale et al., 2019; Dai et al., 2020). The dysregulation of cortisol levels, a symptom associated with stress, also contributes to the disease process’s morbidity, severity, and mortality. This alteration includes numerous oncological diseases such as tumor progression in breast cancer (Sephton et al., 2000; Lillberg et al., 2003; Alejandra Ruiz-Manzano et al., 2019).

Three-dimensional (3D) cultures are a valuable tool to study the functions of cancer genes and pathways in an appropriate polarized context and the interaction between tumor cells. 3D cultures allow for a much better understanding of the pathological characteristics of tumors with the microscope.

The present work evaluates cortisol’s action on mitochondrial activity and cell proliferation and the interaction between Hep-2 tumor cells in 3 D culture, treated with hydrocortisone.

## 2. Materials and Methods

### 2.1. Cell culture

HEp-2 (carcinoma of human larynges) were obtained from the cell bank of Rio de Janeiro and cultured at 37 °C under 5% CO2 in Dulbecco’s Modified Eagle Medium (DMEM, Gibco BRL, Grand Island, NY, USA) supplemented with 10% fetal bovine serum (FBS) (Gibco BRL, Grand Island, NY, USA), and 1% penicillin and streptomycin (Invitrogen Life Technologies, Carlsbad, CA, USA).

### 2.2. Incubation with hydrocortisone

HEp-2 cells were plated (1×10^5^ cells/mL) in 24-well microplates, with MEM culture medium supplemented with 10% fetal bovine serum (SFB) for cell adhesion incubated at a temperature of 37°C and atmosphere 5% CO2, overnight. The next day, the cells were subject to treatment with Hydrocortisone 500 mg, diluted in PBS, for periods of 24 and 48 hours in the following concentrations: 0.5 μM, 1.0 μM, 1.5 μM, 2 μM, and 2.5 μM (Abdanipour et al., 2014).

### 2.3. Mitochondrial Metabolic Activity (MTT assay)

HEp-2 cells submitted to treatment with hydrocortisone in the periods of 24 and 48 hours were washed with PBS three times incubated with MTT (0.5 mg/ml) for 1 hour at 37°C in an atmosphere of 5% CO2. Over the formazan precipitates, the organic solvent DMSO (50 μL) was added to each well. The plate is kept under shaken for 10 minutes to solubilization the formazan crystals and the absorbance reading on a Packard SpectraCount wavelength of 570 nm. The data obtained were plotted in a graph by the GraphPad 6.0 program.

### 2.4. Crystal Violet Assay

Crystal Violet staining is used to infer population density, a clonogenic test. The method is described as colorimetric for determining appropriate cells, as defined by Fernandes et al. (2010). Cristal Violeta, a substance used for the test, crosses cell and nuclear membranes, binding to DNA, RNA, and proteins, identifying viable cells. In its outline, the spectrophotometric quantification of adherent cells indirectly identifies the number of viable cells used in this study to measure the cell lines’ population variation.

The HEp-2, subjected to treatment with cortisol at the mentioned times, had the culture medium removed afterward and incubated with 100 μL of Crystal Violet for 3 minutes at room temperature. After, the plate was washed 2x in tap water by immersion and incubated for 1 hour with 100 μL of DMSO to read on the photometer spectrum at 570 nm.

### 2.5. 3D culture

Cultures were performed using the Bio-AssemblerTM kit designed for 24-well plates (n3D-Biosciences Inc, Houston, TX, USA) (Souza-Araújo et al., 2020). In summary, NanoShuttlesTM was used for incubation. Plates of 24 wells were used, a ratio of 1 μL of NanoShuttlesTM per 20,000 cells, and incubated at 37°C and 5% CO_2_ overnight. The cells were then detached by treating them with 5 mL of trypsin for 5 min and washed by centrifugation (600 g/5 min) with a balanced saline solution (PBS). Cell viability was determined by the trypan blue exclusion method (1% w/v in PBS), and the density was adjusted to 10^6^cells/mL in the medium supplemented with RPMI-1640.

HEp-2 cells conjugated to NanoShuttlesTM were seeded on a 24-well ultralow-attachment plate (ULA, Cellstar® Greiner Bio-one, Kremsmünster, Austria) in 105 cells and a final volume of 400 μL/well.

The 3D culture was obtained by incubating (at 37°C and 5% CO_2_) the plates under the magnetic field, first using a bioprinting unit for three hours, followed by the levitation unit during the entire culture period. This procedure allows for the growth of the cellular spheroid. The 3D culture plate was replenished with fresh medium every two days until using the spheroid.

### 2.4. Immunostaining

After seven days of culture, HEp-2 cell spheroids were divided into two groups: (i) a cells-only control group and (ii) a treatment group, in which the cells were incubated with cortisol (2.5 μM) for 48 hours at 37°C in a 5% atmosphere of CO2. After this, the spheroids were fixed with 4% paraformaldehyde in PBS for 15 minutes at room temperature. The cells were then permeabilized with 0.2% Triton X-100 in PBS for 10 min and then blocked with 1% bovine albumin serum (BSA) in PBS for 30 minutes. After that, the spheroids were incubated with anti-human monoclonal antibody mouse against fibronectin antibody (1:500/1h) and rabbit anti-human monoclonal antibody against FAK (1:500/1h), and then incubated with anti-mouse polyclonal secondary antibody was conjugated to FITC, and goat anti-rabbit polyclonal secondary antibody conjugated to TRITC (all antibodies from Sigma Aldrich, Co). The nuclei were marked with DAPI (4′,6-diamidino-2-phenylindole). Samples were visualized using a fluorescence microscope (DMIL, Leica).

### 2.5. Statistical Analysis

The data are in the form of mean and standard deviation, compared by the two-way ANOVA test and confirmed by the Tukey test. Statistical significance was admitted with P <0.05 with * P <0.05; ** P <0.01 being considered significant. Experiments were performed in three independent replications with n = 8. GraphPad Prism 6® software was used to perform statistical and graphical analyzes.

## Results

The mitochondrial evaluation by MTT aimed to measure possible influences of incubation with hydrocortisone, in concentrations of 0.5, 1.0, 1.5, 2.0, and 2.5 μM, in HEp-2 cell culture, relating possible findings to the population density results which were then analyzed. Figure 1 presents the results of the mitochondrial activity.

**Figure 1.**
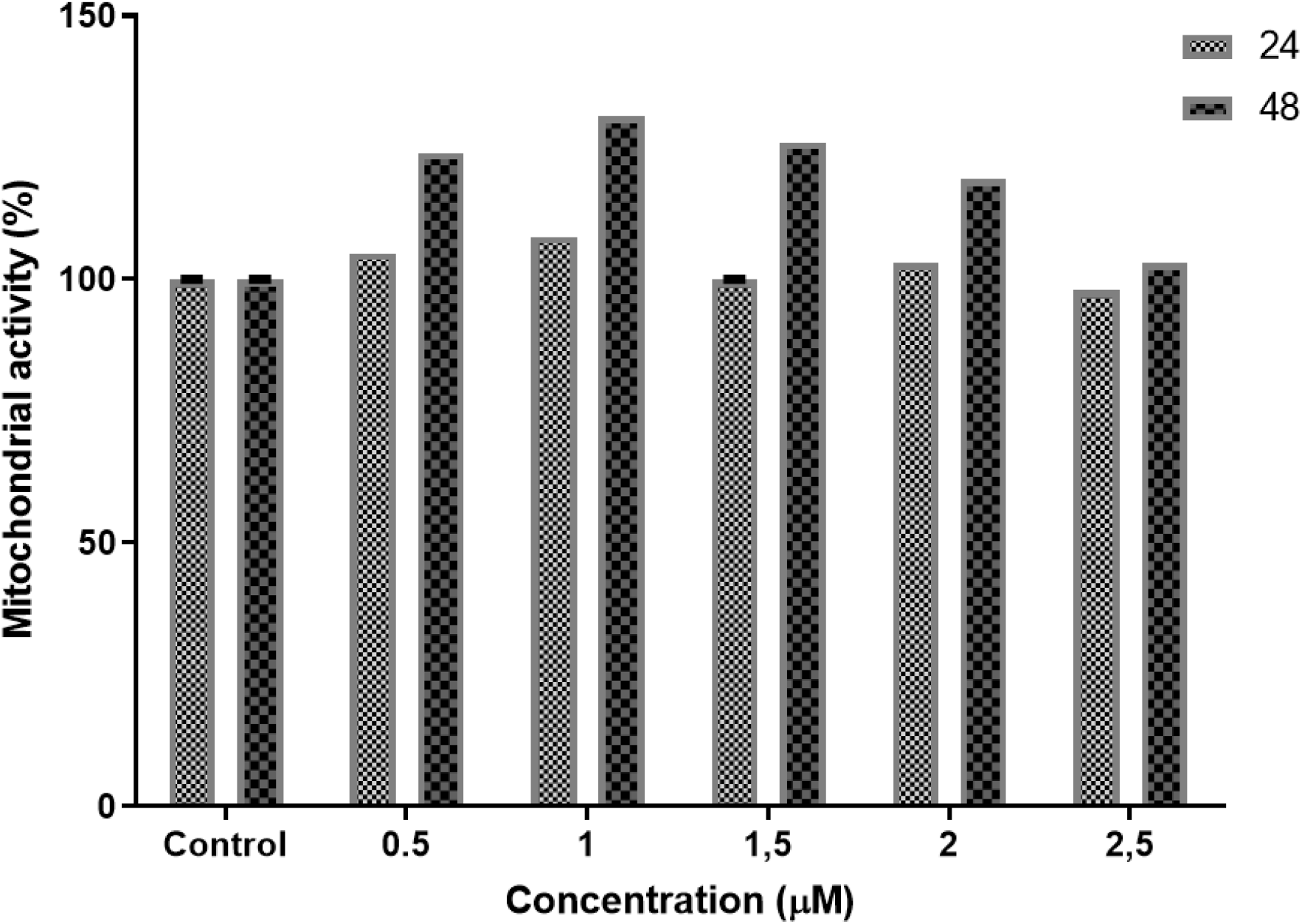
Mitochondrial activity (MTT) assay of the HEp-2 cells. Cellular activity of the HEp-2 cells MTT assay at after 24 and 48, incubation of increasing hydrocortisone concentrations, shows an increase in cell numbers with an increase in time. All values are expressed as mean ± standard error of the mean (SEM) from three different samples

Compared with the control group, the other groups with increasing hydrocortisone concentrations do not show a statistical difference within 24 hours. The comparative analysis of the control group with the other groups in the 24-hour period shows a statistically significant increase in mitochondrial activity, mainly in the control, with concentrations of 0.5 to 2.0 μM (p <0.0001), while the comparison between different concentrations of hydrocortisone results in a significant increase in mitochondrial activity between groups 0.5 vs 2.5 μM (p <0.0001), 1.0 vs 2.0 μM (p <0.0003), 1.0 vs 2.5 μM (p <0.0001), 1.5 vs 2.5 μM (p <0.0001), and 2.0 vs 2.5 μM (p <0.0001). Within 48 hours, a significant increase was observed in all concentrations when compared to the control.

Population growth was assessed using the Crystal Violet test. The data described in Figure 2 reveal that even under exposure to increasing hydrocortisone concentrations, there was no population reduction in the two distinct periods, except for the group incubated at 2.5 μM in the 24 hours, but no significant population reduction was observed.

**Figure 2.**
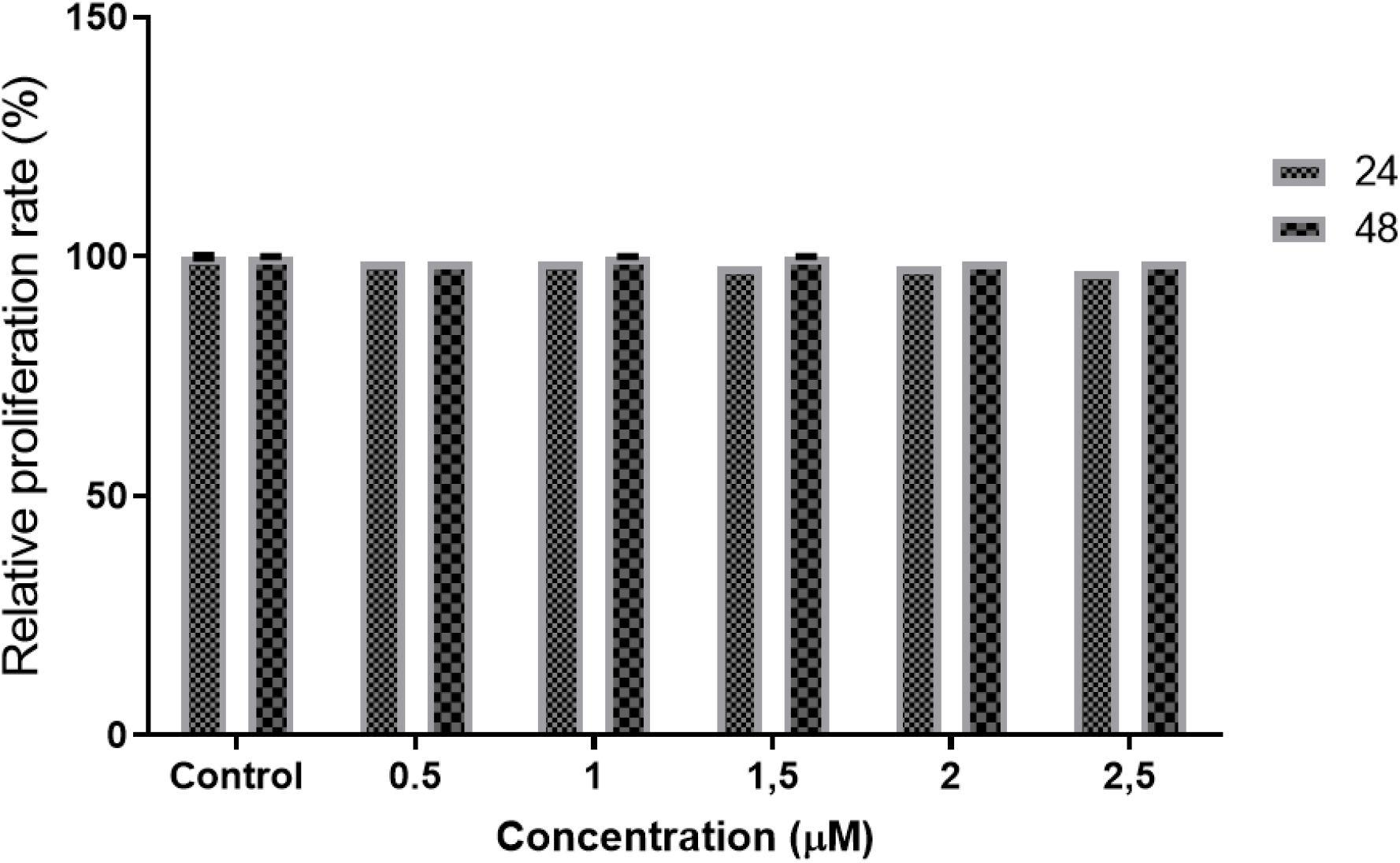
Crystal violet assay. Crystal violet assay of HEp-2 cells, 24 to 48 hours incubation with increasing concentrations of hydrocortisone. All values are expressed as mean ± standard error of the mean (SEM) from three different samples.

The 3D growth assessment of HEp-2 cell culture under the action of hydrocortisone (2.5 μM) after 24 and 48 hours of incubation is shown in Figures 3 and 4.

**Figure 3.**
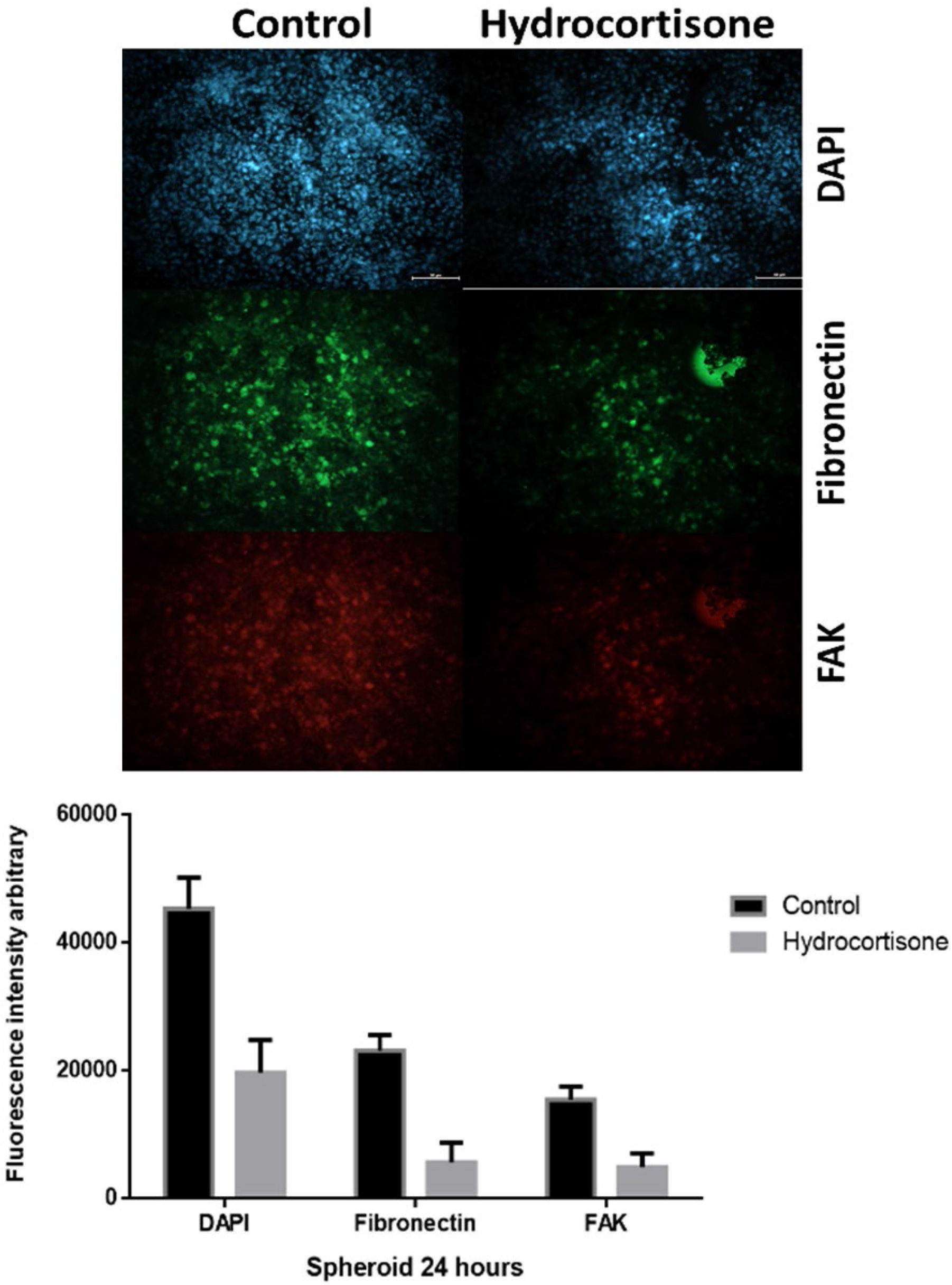
Tumor spheroids immunolabeled for fibronectin and FAK. Cell proliferation and morphology of tumor spheroids at day 7 of culture, and after 24 hours incubation with 2.5 μM of hydrocortisone, A) Photomicrography of spheroids after immunolabeled with ant-fibronectin and anti-FAK, nuclei labeled with DAPI, bar 50 μm. B) Graphic fluorescence intensity for DAPI, fibronectin, and FAK.

**Figure 4.**
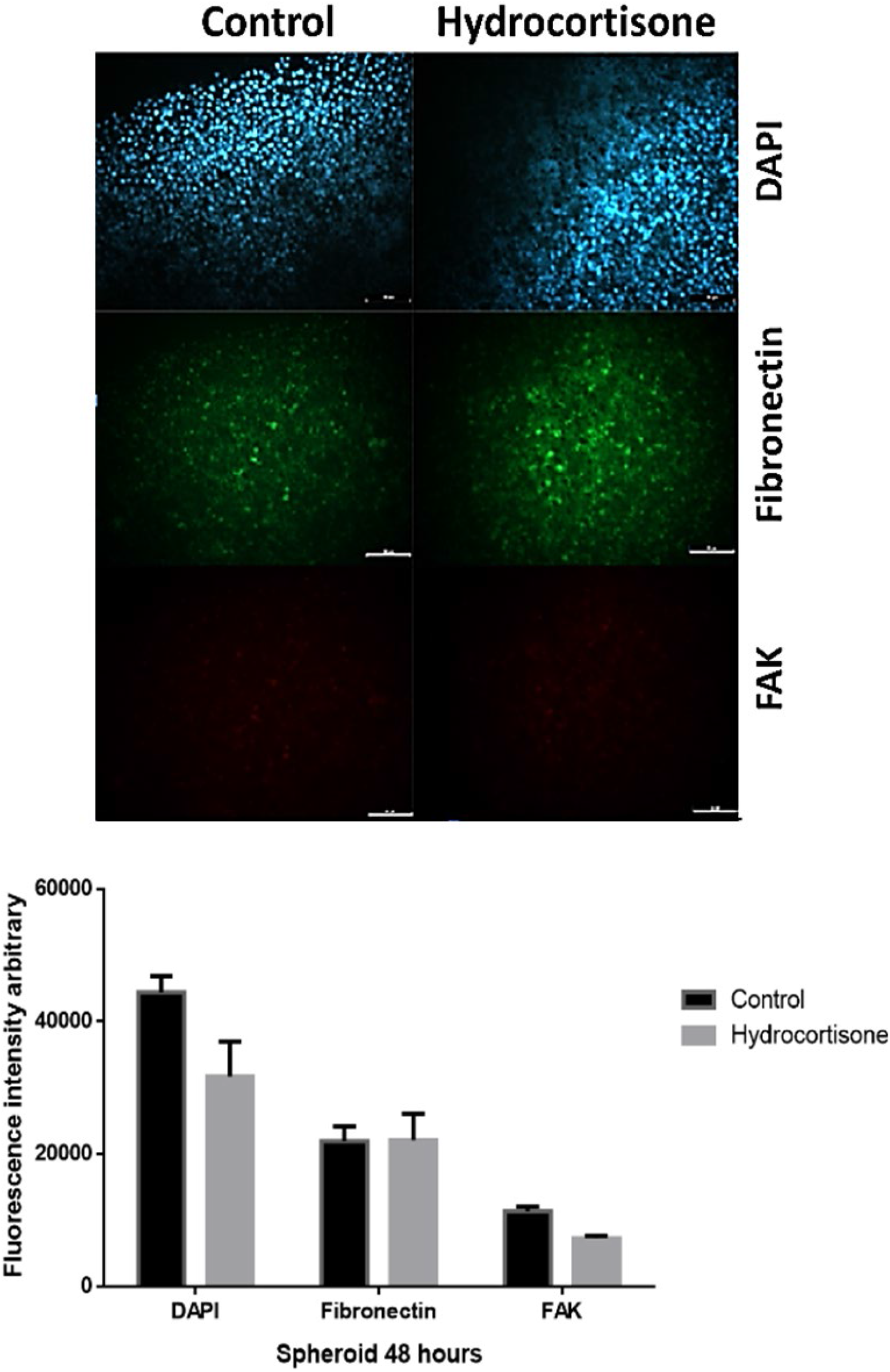
Tumor spheroids immunolabeled for fibronectin and FAK. Cell proliferation and morphology of tumor spheroids at day 7 of culture, and after 48 hours incubation with 2.5 μM of hydrocortisone, A) Photomicrography of spheroids after immunolabeled with ant-fibronectin and anti-FAK, nuclei labeled with DAPI, bar 50 μm. B) Graphic fluorescence intensity for DAPI, fibronectin, and FAK.

In 24 hours, the interaction with hydrocortisone was observed to be interfering with cells’ spheroid formation, showing a significant dispersion of cells marked with DAPI (p = 0.0004). Compared to the control, the immunolabeled for fibronectin exhibited a significant reduction (p = 0.0192) in the staining intensity (Figure 3 b). Within 48 hours, there was no significant change in fluorescence intensity in the immunolabeling observed (Figure 4 b).

Figure 5 illustrates endogenous and exogenous glucocorticoids’ possible mechanisms in tumor cells based on the results obtained.

**Figure 5.**
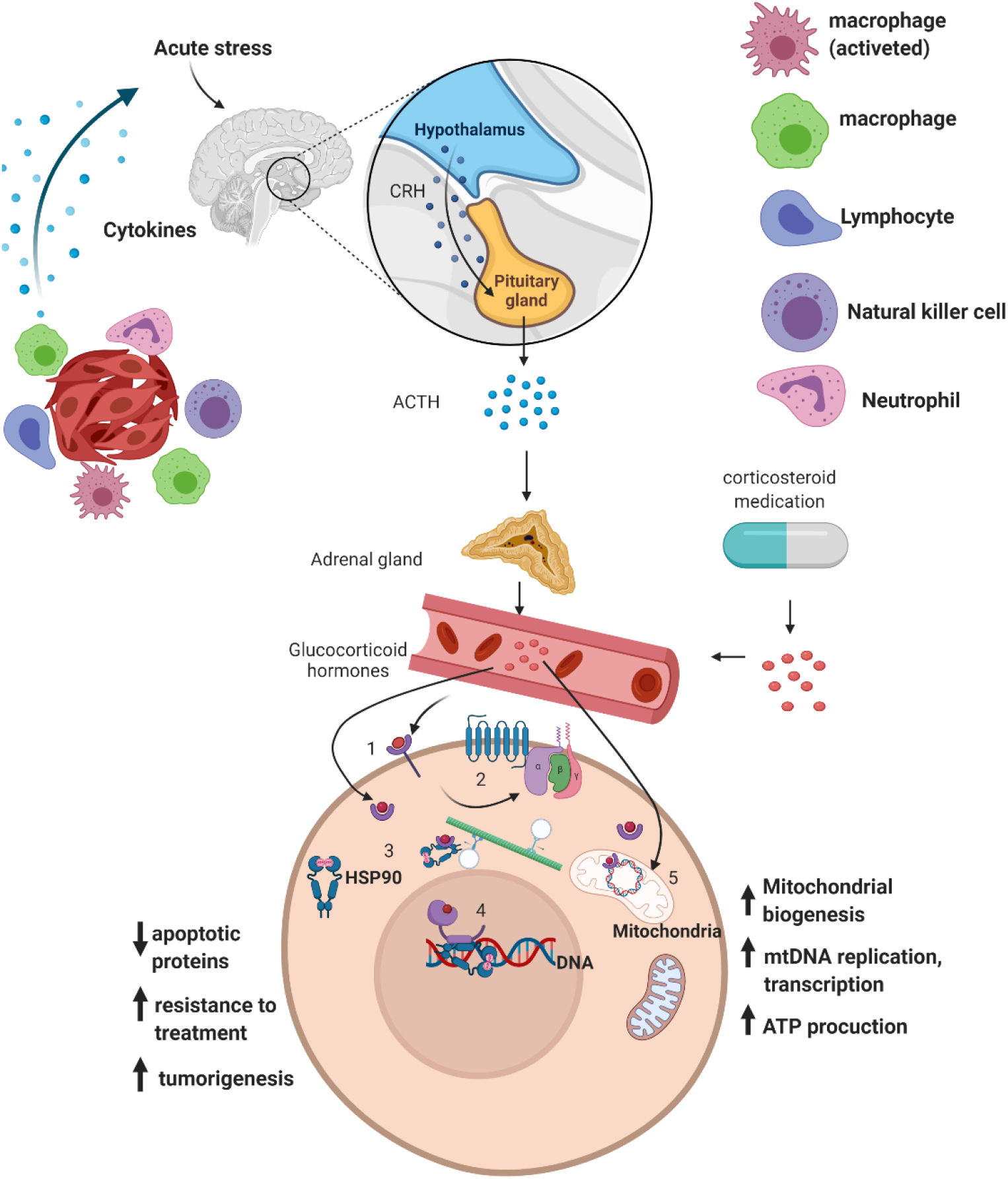
Tumor cells can be subjected to corticoids (glucocorticoids) of endogenous or exogenous origin. The sum of this hormone can cause damage to the tumor cell or act as a trigger for resistance to radiotherapy or chemotherapy treatment. The action of glucocorticoids can be non-genomic or genomic. The action of non-genomic glucocorticoids is mediated by membrane-bound GRs glucocorticoid receptors **(1),** which trigger the activation of kinase signaling pathways. Corticosteroid receptors associated with the membrane can be coupled to the G **(2)** protein, located on the membrane and which, by activating cascades, it triggers signaling by secondary messengers. The genomic action involves the diffusion of glucocorticoids across the cell membrane into the cytoplasm, where they bind to the glucocorticoid receptor (GR). The intracellular GR is linked to stabilizing proteins **(3)**, for example, the heat shock protein 90 (Hsp90). This complex then translates, through interaction with dynein, to the nucleus, where it exerts its effect **(4)**. Another genomic effect can occur in mitochondrial DNA **(5)**, acting as a regulator of functions and energy metabolism in the mitochondria.

## 3. Discussion

The disclosure of cancer diagnosis and treatment is usually a traumatic experience for the patient, causing cortisol release in response to stress associated with physical and mental comorbidities.

To investigate cortisol effects on mitochondrial activity and cell proliferation, we exposed HEp-2 cells to 0.5–2.5 μM cortisol for 24 and 48 hours and subjected them to MTT and crystal violet assays. The 3D culture was also tested to evaluate the extracellular matrix.

The results of the evaluation of the mitochondrial activity show that after 24 hours of exposure to increasing concentrations of hydrocortisone, the cells presented a significant increase at 0.5 and 1.0 μM and a reduction at 2.5 μM. There is a significant increase in mitochondrial activity within 48 hours, indicating intense cellular activity, corroborated by (Bomfim et al. 2018). This mitochondrial activity can be a relation to an increase of glucose, due to the increase in glucocorticoid in the system, corroborate with Seyfried et al., 2010; Manoli et al. 2007, who that report which GC such as dexamethasone, regularly prescribed, increase blood glucose levels, contributing energy to glycolysis-dependent tumors, accelerating the growth of brain tumors, for example.

Another factor that can be influenced the result of the MTT assay is related to the exposure for short periods (24 and 48 hours) to concentrations of GCs is associated with the induction of mitochondrial biogenesis and with the enzymatic activity of selected subunits of the respiratory chain complexes, leading to increased mitochondrial activity, according to Dong et al., 2019.

The crystal violet assay shows DNA duplication. It can be verified that the decrease observed within 24 hours is insignificant and that the duplication of the genetic material and, consequently, the cell proliferation within 48 hours remains like the control. These results are corroborated by Shannon et al. 2015, Dong et al., 2019. GC’s action in DNA involves more transcriptional events associated with the resistance process of tumor or apoptotic mechanism, but this action was associated with the stress type, acute or chronic.

The evaluation of 3D growth HEp-2 cells and treatment with a concentration of 2.5 μM demonstrates that such concentration interferes with the interaction between cells after 24 hours of incubation, presented scaterred cells after incubation with GC. There was no significant change in the interaction and spheroid formation after 48 hours when control compared. Similar results were obtained by Chen et al. 2010 and Alnatsheh, 2019, and the accord (Shannon et al., 2015). The dexamethasone (Dex) treatment also significantly increases cell-ECM adhesion strength and decreases motility in glioblastoma cells.

## 4. Conclusion

Corticosteroids and stress in cancer patients may interfere with cancer treatments because these may cause tumor cells to progress instead of reducing depending on the cell type. Although some cases favored corticosteroid use in cancer patients, a more detailed analysis is necessary before prescribing them. Moreover, it is crucial to assess the patient’s cortisol level before and after treatment as well.

## Acknowledgment

This study had financial support of the Fundação de Amparo à Pesquisa de São Paulo (FAPESP) under Partnership Grant Number 16/17984-1, FINEP/MCTIC – accord n° 01.13.0275.00, and part by Coordenação de Aperfeiçoamento de Pessoal de Nível Superior – Brazil (CAPES) – finance code 001.

